# Identification of non-covalent inhibitors for the atypical peroxiredoxin PRDX5 as a therapeutic strategy in malignant pleural mesothelioma

**DOI:** 10.64898/2026.05.13.724787

**Authors:** Josep Monserrat, Floriane Montanari, Victor Laurent, Pierre-Benoit Ancey, Nicolas Jean, Cécilia Jeannu, Guimin Wang, Genshu You, Qiancheng Shen, Alice Mac Kain, Yacine Bareche, Loïc Herpin, Nadia Jeremiah, Roberta Codato, Alberto Romagnoni, Alex J. Cornish, Ekaterina Rozhavskaya, Lucia Pattarini, Constance Petit, Pierre-Joachim Zindy, Jinal Shukla, Sylvie Gomez, MOSAIC Consortium, Markus Eckstein, Almoatazbellah Youssef, Ulrich Keilholz, Markus Morkel, Krisztian Homicsko, Chiara Saglietti, Leilei Shi, Jian Zhang, Elodie Pronier

**Author notes:** Correspondence should be addressed to E.P.

## Abstract

Malignant pleural mesothelioma (MPM) is an aggressive asbestos-linked cancer with limited therapeutic options and a dismal 5-year survival rate of ∼5%. While aberrant production of reactive oxygen and nitrogen species (ROS/RNS) is a hallmark of MPM, targeted approaches to exploit these redox vulnerabilities remain scarce. Here, using the MOSAIC multimodal cancer patient atlas, we identify Peroxiredoxin 5 (PRDX5) as being significantly upregulated in the epithelioid subtype of MPM. We show that MPM cells exhibit enhanced resistance to nitrosative and oxidative stress compared to healthy mesothelial cells, a phenotype correlated with basal PRDX5 expression. Next, utilising a machine learning guided discovery pipeline, we identified three putative allosteric pockets in PRDX5 and conducted a virtual screen of 3.6 million compounds. High-throughput biochemical validation of 452 candidates yielded 36 non-covalent hits, including sub-micromolar inhibitors. These findings establish PRDX5 as a novel, subtype specific therapeutic target in MPM and provide a chemical framework for the development of next-generation redox-modulating oncology treatments.

## Introduction

Malignant pleural mesothelioma (MPM) is an aggressive malignancy originating from the mesothelial cells of the pleura, with a strong etiological link to asbestos exposure^1–3^. Given the nature of occupational exposure, MPM disproportionately affects men and is characterised by a long latency period, typically 20 to 50 years^1,2,4,5^. Although improved occupational safety and exposure prevention have led to a decline in new cases, MPM continues to represent a significant unmet clinical need.

Current therapeutic options offer limited overall survival and contribute to a stubbornly low 5-year survival rate of approximately 5%^6^. Furthermore, despite distinct histological subtypes of MPM, current treatment regimens lack targeted approaches, often failing to account for these differences^7^. This highlights a critical need for the development of novel, subgroup-specific therapies to improve treatment efficacy and patient outcomes.

Aberrant generation of reactive oxygen species (ROS) and reactive nitrogen species (RNS) is a common feature of various malignancies^8–10^. This heightened production stems from the increased energetic demands of rapidly proliferating tumour cells, and the resulting acidic and hypoxic environments in the tumour microenvironment (TME)^10–12^. To counteract this stress, malignant cells upregulate a suite of protective mechanisms, including reactive species scavengers, DNA-repair machinery and enhanced angiogenesis^13,14^. Consequently, the targeting of these critical stress-coping mechanisms has emerged as a promising therapeutic strategy, aiming to induce cell death by pushing malignant cells beyond their capacity to efficiently manage oxidative and nitric stress^15^.

Given the persistent inflammation, ROS and RNS production associated with asbestos exposure, MPM may be uniquely susceptible to dysregulation of its stress response pathways^16^. This vulnerability has been exploited in MPM preclinical and clinical settings with promising results. For example, thiostrepton, a potent small molecule inhibitor of peroxiredoxin 3 (PRDX3), a key scavenger of mitochondrial-derived ROS, is currently being evaluated in Phase 2 clinical trials, building upon encouraging biomarker data from Phase 1 studies^17^.

Recently, another member of this family, peroxiredoxin 5 (PRDX5), has gained attention as a crucial RNS scavenger that is frequently overexpressed across a wide range of human cancers^18^. PRDX5 has been shown to be functionally important for cancer cell survival, where its pharmacological or genetic inhibition leads to a significant decrease in cell viability, increased apoptosis, and reduced migratory potential^19–21^. Importantly, PRDX5 is structurally unique among the six peroxiredoxin family members, as both active site cysteines (Cp and Cr) react within a single monomer, despite PRDX5 dimer formation for catalytic function. This stands in juxtaposition to the obligate dimeric cysteine-cysteine disulphide bonds between monomers observed in other PRDXs^22^. This structural feature represents a compelling opportunity for the development of highly specific small molecule inhibitors with minimal off-target effects on other peroxiredoxins, decreasing the likelihood of undesired toxic effects.

Here, we investigate the functional role of PRDX5 as a putative therapeutic vulnerability of MPM. We demonstrate that PRDX5 is significantly upregulated in epithelioid MPM, where it mediates a distinctive resistance to oxidative and nitric stress not observed in healthy mesothelial cells. To target this mechanism, we employed machine learning to identify and rank cryptic allosteric pockets in PRDX5 and conducted a virtual screen of non-covalent chemical libraries. Biochemical validation of high-scoring candidates yielded 36 inhibitors across diverse chemical series, representing an 8% hit rate. These findings establish PRDX5 as a tractable therapeutic target in epithelioid MPM and provide a structural framework for the development of small molecule PRDX5 inhibitors.

## Results

### PRDX5 upregulation in epithelioid MPM patients

To investigate the role of PRDX5 in MPM, we first interrogated the MOSAIC cancer dataset^23^, a large multi-indication and multimodal oncology patient data repository. Specifically, we analysed the untreated baseline samples of the MOSAIC MPM cohort using single-cell RNA sequencing for which we had clinical histological annotation (n = 64; Epithelioid = 41, Sarcomatoid = 9, and Biphasic = 14) and spatial transcriptomics (n = 54) to assess *PRDX5* expression in malignant cells. In both analyses we observed an increased number of malignant cells and tumour islets expressing *PRDX5* as compared to *PRDX3*, a clinical-stage related family member primarily responsible for detoxification of ROS (Fig. 1). Furthermore, our analysis revealed that patients of the epithelioid histological subtype exhibited significantly higher PRDX5 expression than those of sarcomatoid or biphasic subtypes (Fig. 1A).

**Figure 1:**
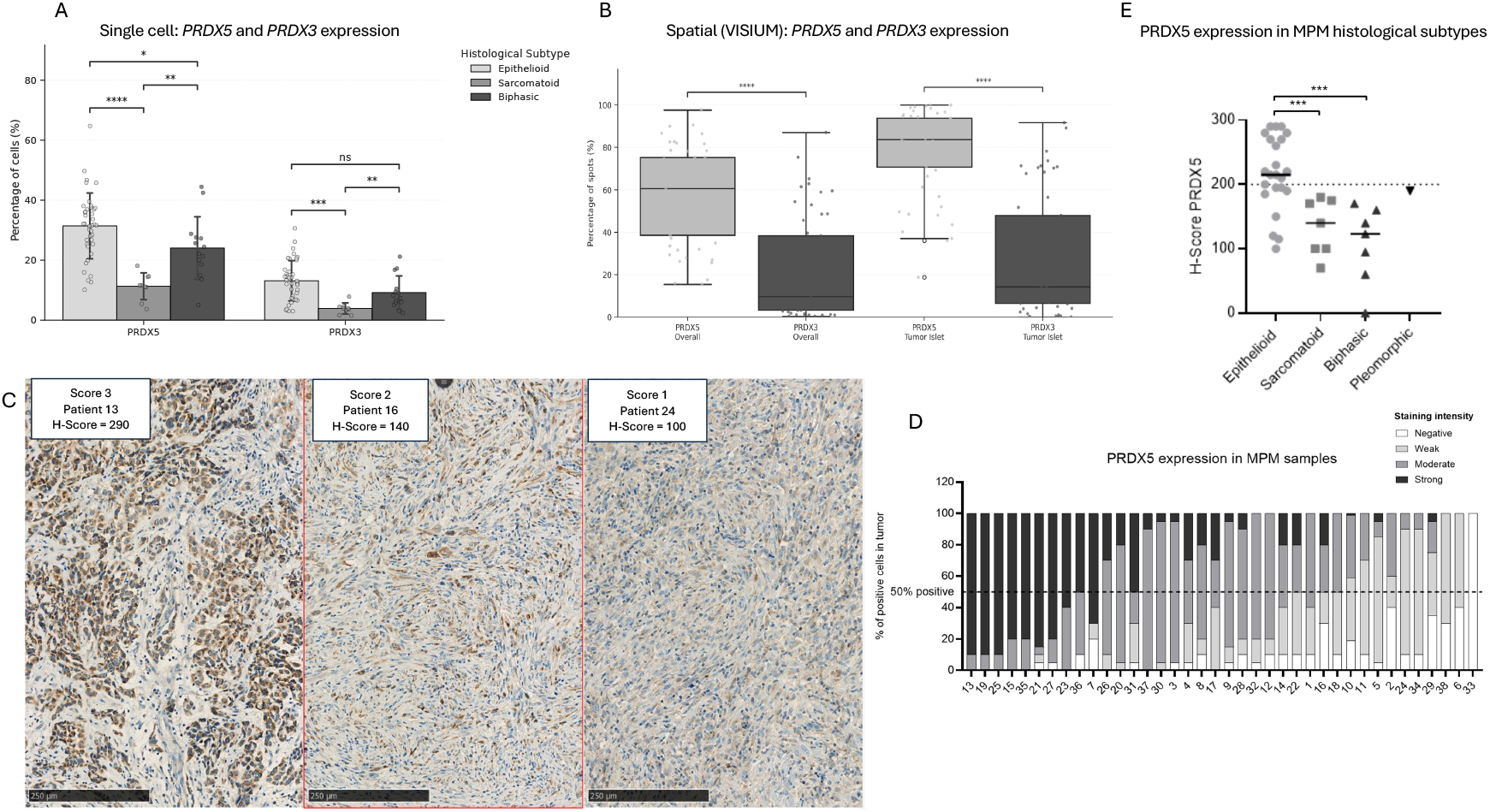
*PRDX5* is highly expressed in epithelioid MPM patients. **A.** Prevalence of *PRDX5* and *PRDX3* expression across mesothelioma histological subtypes. Percentage of malignant mesothelioma cells with detectable expression (logCPM > 0) of *PRDX5* and *PRDX3* across Epithelioid (n = 41), Sarcomatoid (n = 9), and Biphasic (n = 14) histological subtypes. Data are derived from single-nuclei transcriptomics of baseline mesothelioma samples from the MOSAIC dataset; bars represent mean values across samples and error bars indicate the standard deviation. Significance was determined by a two-sided Mann-Whitney test (****P < 0.0001, ***P < 0.001, **P < 0.01, *P < 0.05, ns = non-significant. **B.** Differential spatial prevalence of *PRDX5* and *PRDX3* expression within the overall slide or the tumour islets. Box plots represent the percentage of Visium spatial transcriptomics spots with detectable expression (logCPM > 0) of *PRDX5* and *PRDX3* when comparing overall tissue sections to morphologically defined Tumour Islets (n = 54 samples) from the MOSAIC dataset. Significance was determined by a two-sided Wilcoxon signed-rank test followed by Benjamini–Hochberg FDR adjustment (****P < 0.0001); centre lines indicate medians, boxes indicate the interquartile range (IQR), and whiskers extend to 1.5xIQR. **C.** Representative images from three patients with differential PRDX5 expression from independent commercial patient cohort obtained from CRB (see Methods). Patients 13 (left) and 24 (right) are epithelioid and patient 16 is biphasic (middle). **D.** H-score components depicting the distribution of PRDX5 expression across all screened mesothelioma patients from the CRB cohort (n = 38). Black indicates strong staining (+++), dark grey moderate (++), light grey weak (+), and white negative staining. **E.** Summary of PRDX5 H-scores across mesothelioma patients from the CRB validation cohort (n = 38, see Methods). The H-score for each patient was calculated using the standard formula: 3×(percentage of strongly stained cells, +++), 2×(percentage of moderately stained cells, ++), and 1×(percentage of weakly stained cells, +). This scoring system yields values ranging from 0 to 300, integrating both staining intensity and the proportion of positive cells to reflect overall PRDX5 expression. Patients are grouped according to the three major mesothelioma subtypes. Note one patient had mixed pleiomorphic histology.

To validate the findings from the MOSAIC consortium, we next quantified PRDX5 expression in primary tumour samples from an independent validation cohort of 38 additional MPM patients (CRB, Centre de Ressources Biologiques, France, see Methods). Our results showed that PRDX5 was expressed in >50% of tumour cells in 97% (37/38) of the patients, with moderate to strong intensity detected in as many as 73% (28/38) of patients (Fig. 1C, D). Importantly, these results corroborated our initial observations, confirming that epithelioid MPM patients presented higher PRDX5 expression (65%, 15/23 patients with H-score >200) compared to sarcomatoid and biphasic patients (Fig. 1E). These data indicate that epithelioid patients, who constitute up to 60% of all MPM cases, may specifically benefit from therapeutic modulation of PRDX5.

### PRDX5-Mediated Detoxification of Reactive Species in MPM

The TME significantly influences local signalling and tumour progression. Among many processes in the TME, hypoxia, a common consequence of uncontrolled cell growth, heavily promotes the generation of ROS and RNS^24,25^.

Following our observation of increased *PRDX5* expression in MPM patients, we hypothesised that malignant cells may leverage these abnormally high levels of PRDX5-mediated detoxification to survive in a hypoxic and ROS/RNS-rich TME. To test this hypothesis, we first classified single cells from our MOSAIC MPM cohorts into *PRDX5* expressors or not expressors and within each group assessed the expression of validated hypoxic and ROS/RNS detoxification gene signatures from the Reactome and Gene Ontology (GO) databases (Reactome Cellular response to hypoxia; Reactome Detoxification of Reactive Oxygen Species; GO Biological Process: RNS metabolic process). This analysis revealed that malignant cells with non-zero *PRDX5* expression were associated with significantly higher activation of these pathways, suggesting that MPM malignant cells upregulate *PRDX5* expression to specifically detoxify hypoxia-related reactive species (Fig. 2A-C). In line with these reactive species originating in the core hypoxic TME, we observed a higher prevalence of these respective signatures in the tumour centres versus the borders in our spatial transcriptomics analysis (Fig. 2D-E).

**Figure 2:**
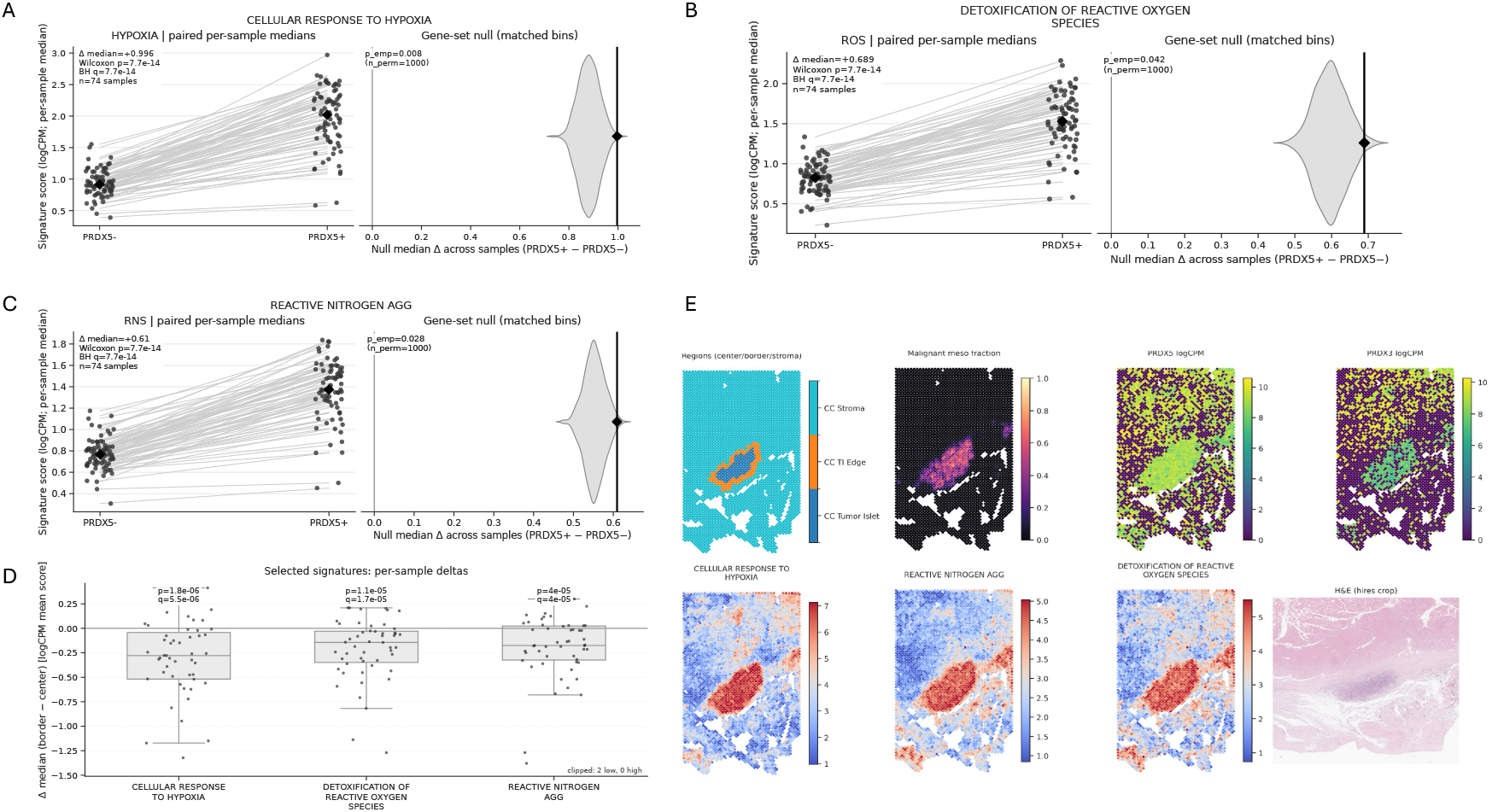
*PRDX5* expression levels correlate with detoxification signatures and the tumour core. **A.** Hypoxia signature expression is enriched in PRDX5^+^ malignant single cells from the MOSAIC MPM cohort. Left, paired per-sample median hypoxia signature scores in PRDX5^−^ (logCPM = 0) versus PRDX5^+^ (logCPM > 0) malignant cells (n = 74 samples; two-sided Wilcoxon signed-rank test; q value adjusted for FDR). Right, specificity of the observed median difference (black diamond) compared to an empirical null distribution generated from 1,000 expression-matched random gene sets (p_emp_ = 0.008). **B.** Detoxification of ROS signature expression is enriched in PRDX5^+^ malignant single cells from the MOSAIC MPM cohort. Left, paired per-sample median hypoxia signature scores in PRDX5^−^ (logCPM = 0) versus PRDX5^+^ (logCPM > 0) malignant cells (n = 74 samples; two-sided Wilcoxon signed-rank test; q value adjusted for FDR). Right, specificity of the observed median difference (black diamond) compared to an empirical null distribution generated from 1,000 expression-matched random gene sets (p_emp_ = 0.042). **C.** RNS signature expression is enriched in PRDX5^+^ malignant single cells from the MOSAIC MPM cohort. Left, paired per-sample median hypoxia signature scores in PRDX5^−^ (logCPM = 0) versus PRDX5^+^ (logCPM > 0) malignant cells (n = 74 samples; two-sided Wilcoxon signed-rank test; q value adjusted for FDR). Right, specificity of the observed median difference (black diamond) compared to an empirical null distribution generated from 1,000 expression-matched random gene sets (p_emp_ = 0.028). **D.** Differential activity of oxidative and nitric stress and hypoxia signatures between tumour regions. Box plots show the paired difference (delta) in median signature scores between the tumour island edge (border) and the tumor islet (center) across the VISIUM spatial transcriptomics cohort (n = 54 samples). Centre lines represent medians, boxes indicate the interquartile range (IQR), and whiskers extend to 1.5xIQR; individual points represent per-sample deltas. Significance was assessed using a two-sided Wilcoxon signed-rank test with Benjamini–Hochberg FDR adjustment (q values shown). **E.** Representative VISIUM spatial transcriptomic images for one mesothelioma MOSAIC patient showing the tumour segmentation, malignant cell fraction, expression of *PRDX5* and *PRDX3*, expression of analysed molecular signatures and H&E staining.

To experimentally validate these observations, we treated three MPM malignant cell lines (IST-MES-1 [epithelioid], IST-MES-2 [epithelioid], and MSTO-211H [biphasic]) and one healthy mesothelial control line (Met-5A) with a range of peroxynitrite and hydrogen peroxide (H_2_O_2_) concentrations to simulate RNS-induced and ROS-induced cellular stress, respectively. Cell viability was assessed as a direct measure of stress-induced cell death. Consistent with our *in silico* findings, all three MPM lines demonstrated a higher resistance to RNS and ROS compared to the healthy mesothelial line (peroxynitrate IC_50_ = 2.21µM, 2.84µM, 2.59µM, and 1.58µM; and H_2_O_2_ IC_50_ = 83.94µM, 360.9µM, 82.31µM and 35.13 µM for IST-MES-1, IST-MES-2, MSTO-211H, and Met-5A, respectively, Fig. 3A). Importantly, this enhanced resistance directly correlated with higher basal expression of PRDX5 in the three MPM lines, while lower levels were observed in healthy Met-5A cells (Fig. 3B). This suggests that PRDX5-mediated detoxification may account for the increased resistance to cell death observed in the malignant lines. Notably, the levels of PRDX3 were comparable among all MPM and healthy cell lines, with higher expression only in IST-MES-2 (Fig. 3B). The observation that IST-MES-2 exhibits a stronger resistance to H_2_O_2_ treatment is supported by the reported higher affinity for ROS detoxification by PRDX3 over PRDX5 and the slightly higher levels of PRDX3 expression^22,26^. In agreement with previous studies^26,27^, our data indicate that PRDX5 plays an important role in ROS and RNS detoxification.

**Figure 3:**
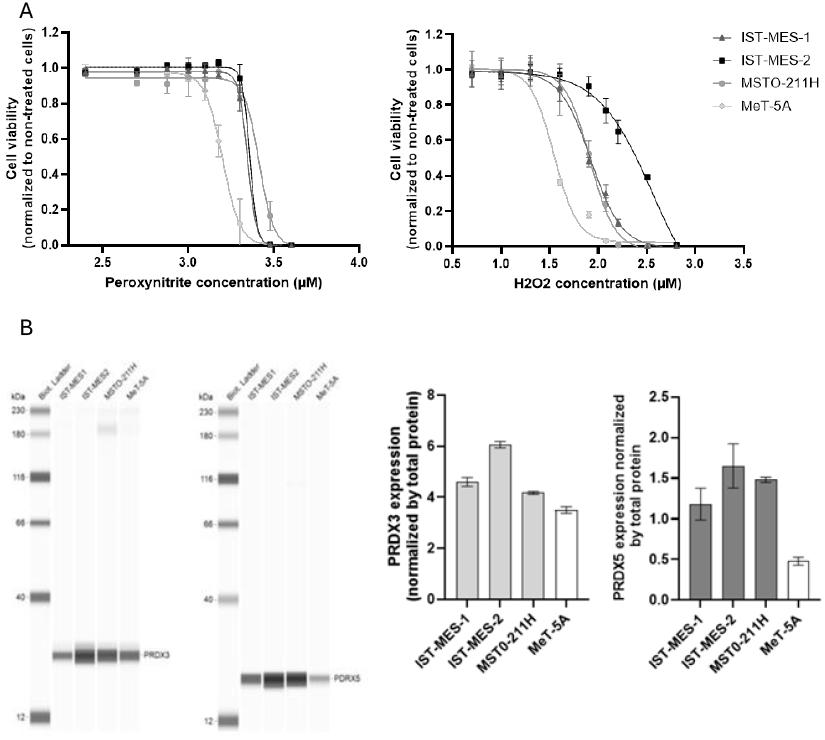
Sensitivity to ROS and RNS is mediated by PRDX5 and PRDX3 in MPM cell lines. **A.** IST-MES-1, IST-MES-2, MSTO-211H and MeT-5A cells were incubated for 5 days in presence of exogenous Peroxynitrite (left panel) or H_2_O_2_ (right panel) at different concentrations. Cell viability was measured using CellTiter assay (luminescence). **B.** Digital representation of data obtained for PRDX3 and PRDX5 detection (capillary western blot) in cell line panel (left panel). Bar plots showing data from one individual run (2 replicates per condition, SD value, right panel).

Based on our observations, we hypothesised that specific inhibition of PRDX5 in epithelioid MPM patients may lead to a positive therapeutic effect. In light of the lack of specific PRDX5 inhibitors, we established a small molecule biochemical screening assay to identify compounds that specifically inhibit PRDX5^22^. However, due to the inherent instability of peroxynitrite, we developed the assay using the more stable H_2_O_2_ as a substrate, given PRDX5 also reduces ROS, albeit at a lower efficiency than RNS (Suppl. Fig. S1)^26,27^.

### Structural biology of PRDX5

Currently, 12 crystal structures of human PRDX5 are available in the Protein Data Bank (PDB). In this study, seven structures were selected based on the following criteria: (i) the protein must be in its reduced state to facilitate inhibitor binding; (ii) the protein must be wild type with no mutations; and (iii) the structural resolution must be better than 2.5 Å. The biological assemblies of these seven structures were employed. Specifically, 1H4O exists as a monomer, while the remaining six structures are dimers. Except for the apo structure 2VL3, the other six structures are in complex with fragment-like ligands bound at the substrate-binding site (the catalytic site containing C_p_: Cys47) (Tab. 1, Fig. 4).

**Table 1.**
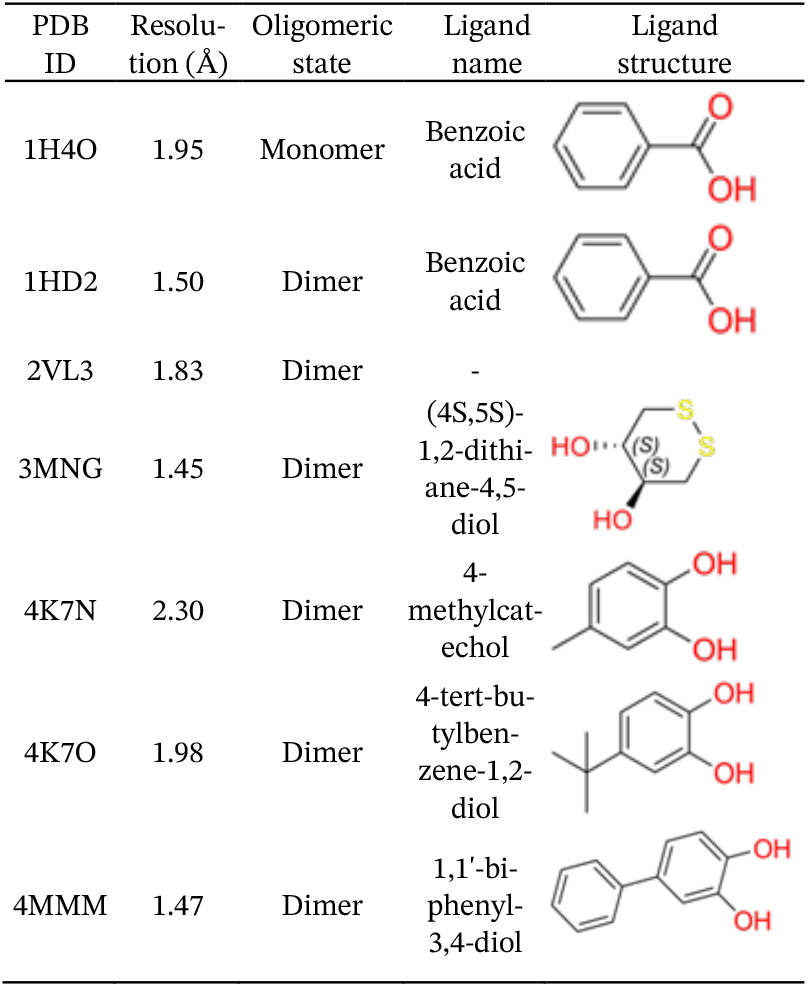
Selected human PRDX5 structures.

| PDB ID | Resolution (Å) | Oligomeric state | Ligand name | Ligand structure |
| --- | --- | --- | --- | --- |
| 1H4O | 1.95 | Monomer | Benzoic acid |  |
| 1HD2 | 1.50 | Dimer | Benzoic acid |  |
| 2VL3 | 1.83 | Dimer | - |  |
| 3MNG | 1.45 | Dimer | (4S,5S)-1,2-dithiane-4,5-diol |  |
| 4K7N | 2.30 | Dimer | 4-methylcatechol |  |
| 4K7O | 1.98 | Dimer | 4-tert-butylbenzene-1,2-diol |  |
| 4MMM | 1.47 | Dimer | 1,1'-biphenyl-3,4-diol |  |

**Figure 4.**
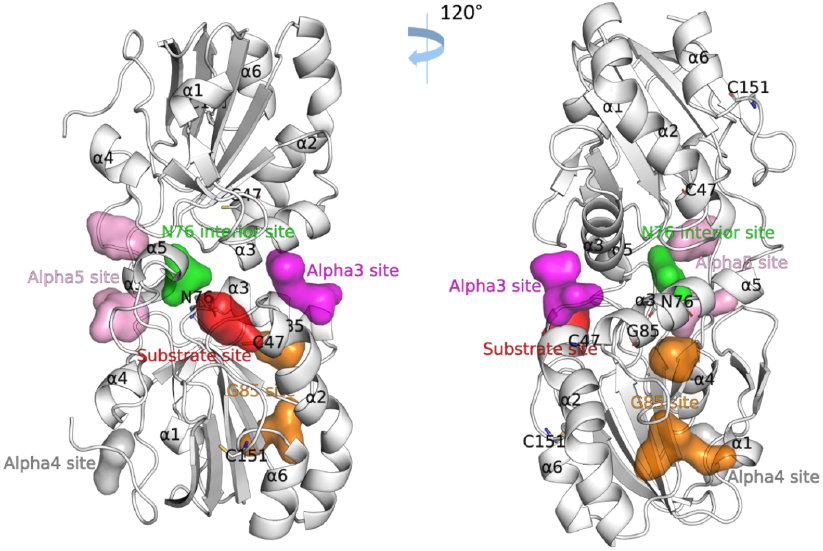
Predicted allosteric sites of PRDX5 as in PDB 4MMM. The protein is depicted as a white cartoon, with the helices labelled. The allosteric sites are represented as a colored surface. The positions of the residues Cys47, Cys151, N76 and G85 are indicated, with their side chains shown as sticks.

All seven PRDX5 structures were prepared and aligned for comparative analysis. The structures exhibited high conformational similarity, with pairwise root-mean-square deviations (RMSDs) of no more than 1.00 Å (Suppl. Fig. S2).

### Identification of putative allosteric pockets on PRDX5

In addition to the substrate-binding site, we aimed to identify potential allosteric sites suitable for non-covalent inhibitor binding. To this end, we employed two internal computational tools from the AlloStar™ platform: AlloDeep and AlloTango. AlloDeep quantifies the allosteric potential of protein residues by utilising elastic networks and heat conduction to capture the regulatory relationships between distant sites. Complementarily, AlloTango evaluates the functional coupling between orthosteric and allosteric sites through molecular dynamics (MD) simulations using the APSF force field^28^. Ultimately, five candidate allosteric sites were identified (Table 2 and Fig. 4).

**Table 2.**
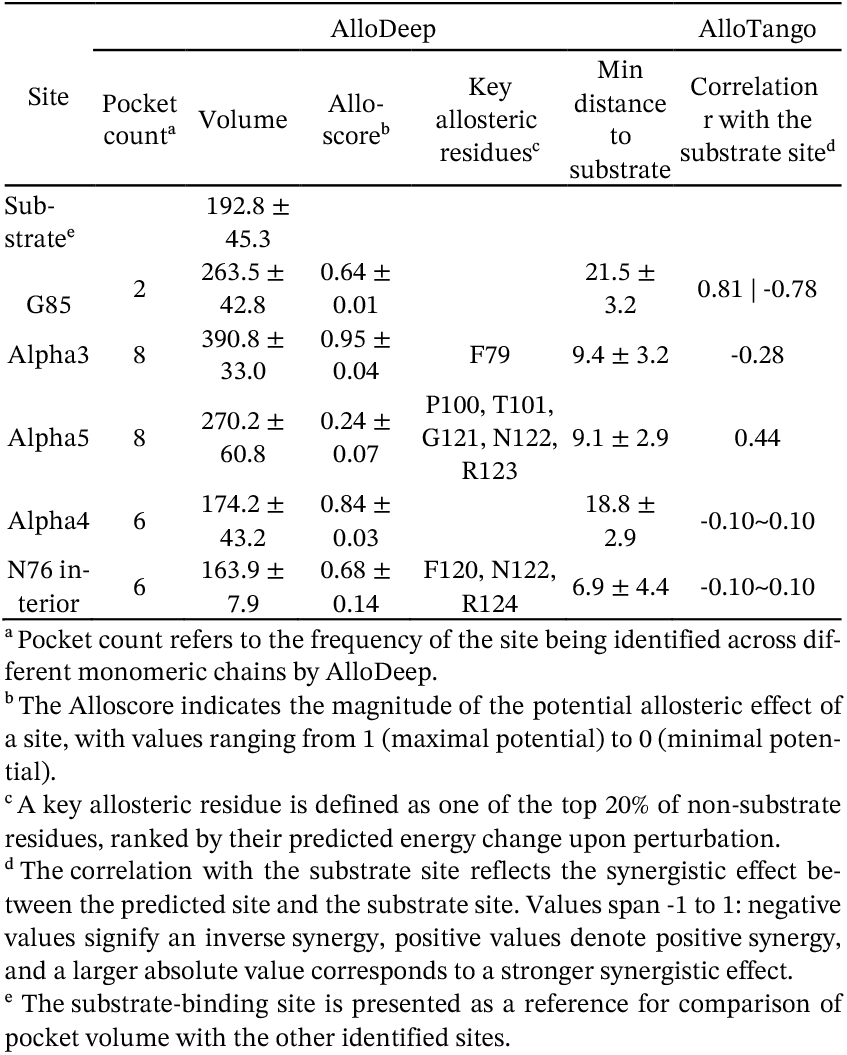
Predicted properties of all the identified sites.

The G85 site is located at the junction of the α2 helix, α3 helix, and β5 sheet. Although initially detected as two adjacent pockets bisected by the α3-β5 loop, they were merged into a single site due to their spatial proximity and limited individual volumes. This site was named after the residue Gly85 on the α3 helix, which exhibits a profound allosteric effect. According to AlloTango, the G85 site shows a strong positive correlation (r = 0.81) with the substrate site on the same chain and a significant negative correlation (r = -0.78) with the substrate site on the symmetry-related chain of the dimer. The high absolute correlation coefficients (|r| > 0.5) indicate that the G85 site possesses the highest synergy with the catalytic centre among all candidates.

The alpha3 site is situated at the dimer interface, flanked by the α3 helix of one chain and the N-terminal end of the α2 helix from the opposing chain. The PRDX5 homodimer adopts a conformation belonging to the C2 point group, characterised by a two-fold rotational axis passing through the interface. Consequently, two symmetry-equivalent alpha3 sites are present. As detailed in Table 2, the alpha3 site yielded the highest Alloscore, suggesting maximum potential for allosteric regulation.

Two additional sites, the alpha4 site (adjacent to the α4 helix and the N-terminal loop) and the N76 interior site (near residue Asn76 within the interface), were also identified. However, both were excluded from further analysis due to their restricted volumes and lack of functional correlation with the substrate site. In summary, the G85, alpha3, and alpha5 sites were selected as the target pockets for subsequent virtual screening.

The alpha5 site is also located at the dimer interface, encapsulated by two α5 helices. Similarly to the alpha3 site, two symmetric alpha5 sites were identified and treated as a unified target for screening. Despite having a lower Alloscore, the alpha5 site contains a high density of key allosteric residues that exhibit the most significant allosteric effects among all the non-substrate residues.

### Virtual screening and non-covalent hit selection

The Enamine Screening Collections (December 2024), comprising approximately 4.4 million compounds, was employed as the initial chemical library. To ensure lead-like quality and reduce false positives, the library was filtered using Pan-Assay Interference Compounds (PAINS) and Rapid Elimination of Swill (REOS) rules. Additionally, a molecular weight (MW) filter was applied to retain molecules between 150 and 500 Da. The resulting filtered library of ∼3.6 million compounds was prepared for docking.

A two-step selection protocol was implemented to identify the most suitable PDB structures for each target site. First, structures were filtered based on the geometric accessibility of the pockets identified by AlloDeep (Suppl. Fig. 3); for the substrate site, all available structures were retained. Second, a pilot docking simulation was conducted using a representative subset of 10,000 compounds randomly sampled from the prepared library with smina^29^. PDB structures were ranked based on two metrics: (i) the median docking score of the entire sample set, and (ii) the average score of the top 100 compounds (Suppl. Fig. 3). Structures demonstrating superior enrichment potential (typically the top three performers) were selected for the final production screening. As detailed in Table 3, this process yielded an ensemble of four PDB structures for the substrate site, two for the G85 site, and three each for the alpha3 and alpha5 sites.

**Table 3.** Selected PDB structures for screening.

| PDB ID | Substrate site | G85 site | Alpha3 site | Alpha5 site |
| --- | --- | --- | --- | --- |
| 1H4O |  | ✓ |  |  |
| 1HD2 | ✓ |  | ✓ | ✓ |
| 2VL3 |  |  | ✓ |  |
| 3MNG | ✓ |  | ✓ |  |
| 4K7N | ✓ |  |  |  |
| 4K7O |  |  |  | ✓ |
| 4MM | ✓ | ✓ |  | ✓ |

To optimise computational efficiency while maintaining chemical space coverage, a two-round screening strategy guided by compound clustering was implemented (Fig. 5). Initially, the entire library was partitioned into 100,000 clusters using the K-means algorithm. In the first round, representative compounds were sampled from each cluster: one compound was selected for clusters containing fewer than 100 members, while for larger clusters, representatives were sampled at a 1:100 ratio. Following the initial docking, the second round focused on the full expansion of clusters associated with the top 1,000 leads identified in the first stage.

**Figure 5.**
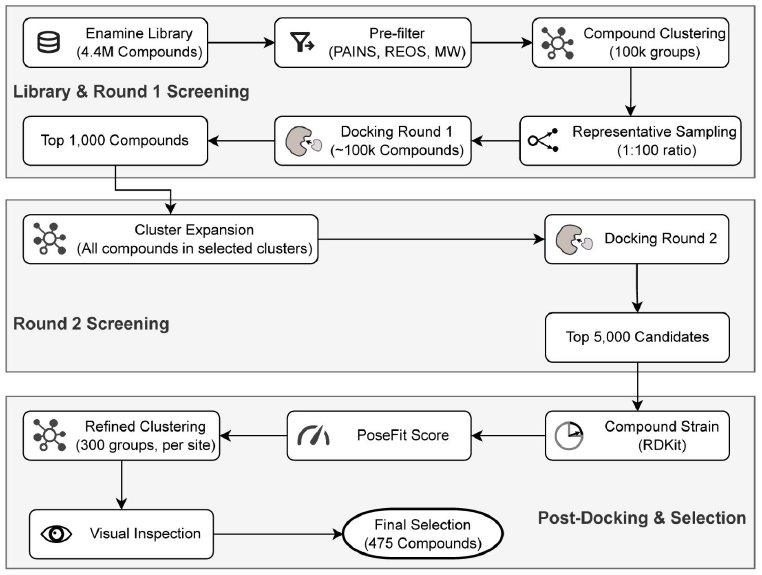
The compound virtual screening workflow.

Post-docking, the top 5,000 candidates underwent rigorous filtering based on two primary criteria: geometric and interaction plausibility. First, the internal strain energy of each docked pose was calculated and compounds exceeding a threshold of 12 kcal/mol were discarded to ensure structural feasibility. Second, pose validity was assessed using PoseFit, a proprietary tool that quantifies the geometric rationality of binding modes. PoseFit generates a score (0 to 1) by comparing the interaction fingerprints of the docked poses with conserved fragment-binding patterns derived from the PDB database. Higher scores indicate superior alignment with experimentally validated interaction motifs.

This workflow was applied across all 12 PDB structures targeting the substrate-binding site and three identified sites (Fig. 5). For each site, the top 300 compounds from their respective PDB ensembles were aggregated and further clustered into 300 groups to ensure chemical diversity. Finally, the binding poses within each cluster were subjected to manual visual inspection. Ultimately, 475 high-priority compounds were selected for experimental validation, of which 452 were successfully procured and tested (Table 4).

**Table 4.** Selected compounds from virtual screening and the hit compounds.

|  | Substrate | G85 | Alpha5 | Alpha3 | Total |
| --- | --- | --- | --- | --- | --- |
| Selected compounds | 143 | 109 | 127 | 96 | 475 |
| Tested compounds | 131 | 105 | 124 | 92 | 452 |
| Active compounds | 4 | 4 | 4 | 1 | 13 |
| Best AC<sub>50</sub> (μM) | 3.89 | 4.46 | 12.01 | 9.81 | 3.89 |
| Average AC<sub>50</sub> (μM) | 12.04 ±4.08 | 10.08 ±2.81 | 15.05 ±2.26 | 9.81 | 12.19 ±3.08 |
| Median AC<sub>50</sub> (μM) | 12.55 | 11.49 | 14.32 | 9.81 | 12.42 |
| Active & Weakly active compounds<sup>a</sup> | 13 | 9 | 11 | 3 | 36 |

<sup>a</sup> The AC<sub>50</sub> of Weakly active compounds is > 30 μM

### Biochemistry screening assay results

All prioritised 452 compounds were tested *in vitro* in a biochemical functional assay^30^ (Thioredoxin assay, see Methods for details), along with DMSO and auranofin as negative and positive controls, respectively. When possible, IC_50_ values were derived. In total, 36 compounds showed inhibition activity in the thioredoxin assay. These active compounds are divided into 3 categories based on data mode curve, activity and potency: 3 compounds have increasing data mode curve and IC_50_ are below 10 µM; 10 compounds have increasing curve and IC_50_ are above 10 µM and still within the measured concentration range; 23 compounds have increasing weakly active curve and a IC_50_ above 30 µM (Table 4 and Fig. 6). All the other compounds remained inactive. All data including chemical structures in a computer-readable format are available in the supplementary information.

**Figure 6.**
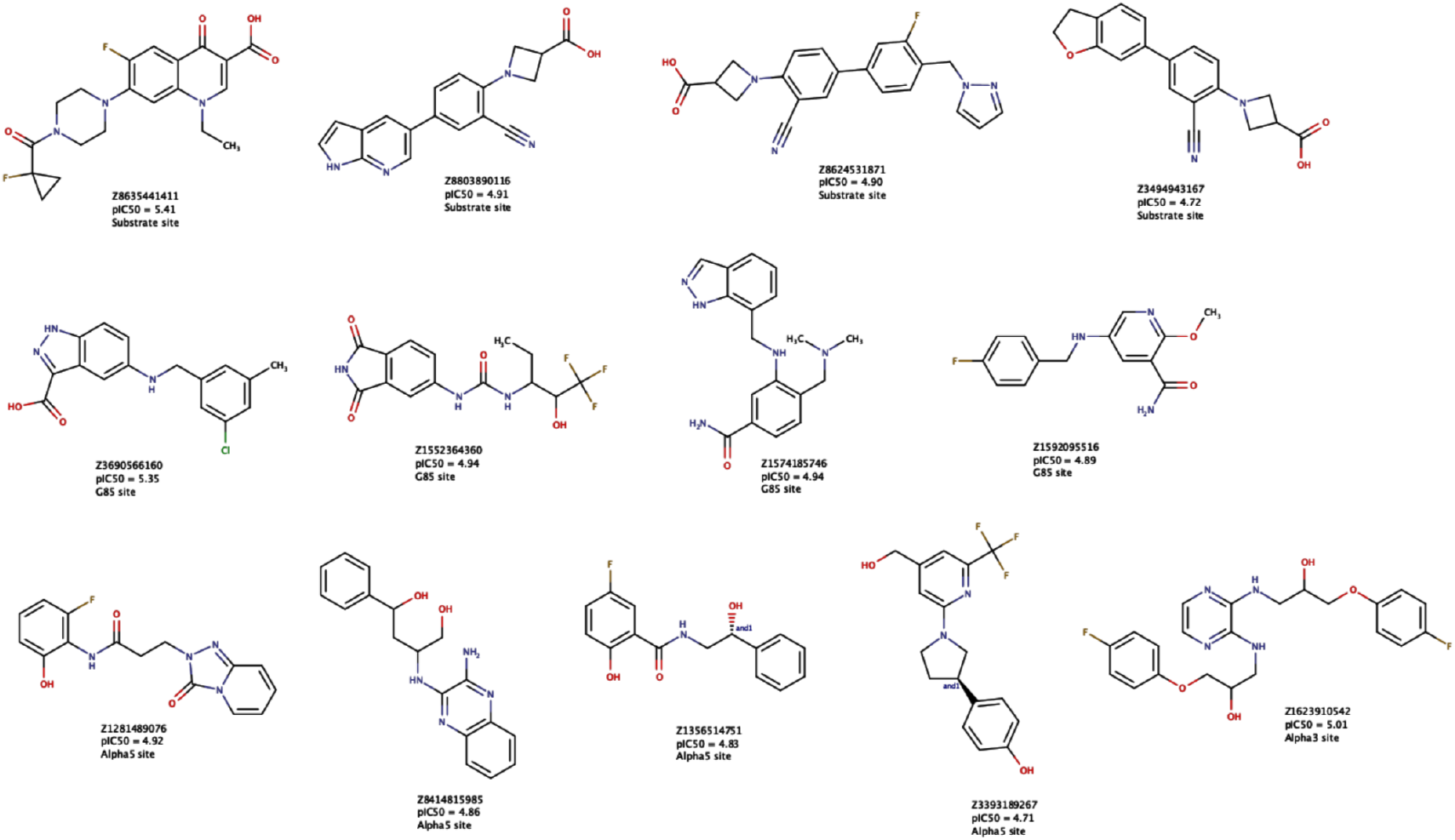
Chemical structures, potency and target site in PRDX5 of the 13 identified hits.

The success of our machine learning screening is evidenced by an 8% hit rate, significantly higher than typical random high-throughput screens (∼1%). By targeting three predicted allosteric sites and the substrate site, we identified 13 high potency and 23 weakly active inhibitors. The G85 and Alpha3 sites, in particular, yielded leads with AC_50_ values as low as 4.46 µM and 9.81 µM, respectively, validating the functional relevance of these previously uncharacterised pockets. With several series exhibiting lead-like properties and medicinal chemistry tractability, including an azetidine moiety, these results confirm that the atypical structure of PRDX5 is not only theoretically druggable but responsive to non-covalent, small molecule inhibition. These 36 hits represent a significant expansion of the PRDX5 chemical space and provide the necessary tools to validate the therapeutic window of PRDX5 inhibition in MPM.

## Discussion

The clinical management of malignant pleural mesothelioma remains a challenge, largely due to the lack of targeted therapeutic approaches and a landscape characterised by a low survival rate of ∼5%^1,6^. Our study sheds light into this unmet need by nominating PRDX5 as a novel, targetable metabolic vulnerability specifically enriched in the epithelioid subtype. By integrating spatial transcriptomics with AI-driven drug discovery, we have identified a first-in-class series of non-covalent inhibitors that exploit the unique structural features of this atypical antioxidant enzyme.

A defining feature of MPM pathogenesis is its etiological link to asbestos fibres, which trigger a cycle of chronic inflammation and the persistent generation of ROS and RNS^10,16^. While previous redox strategies have focused on the mitochondrial scavenger PRDX3, our analysis of the MOSAIC patient atlas reveals that PRDX5 is more broadly expressed in malignant islets and significantly upregulated in the epithelioid population. Unlike the other five peroxiredoxin family members, which function as obligate homodimers with inter-subunit disulfide bonds, PRDX5 operates via an intra-monomer catalytic cycle^22,27^. This unique monomeric architecture provides a critical structural “hook” for achieving isoform selectivity, a long-standing hurdle in the development of peroxiredoxin inhibitors, thereby minimising potential off-target toxicity in healthy tissues.

Our identification of 36 non-covalent hits represents a strategic departure from traditional covalent redox inhibitors, which often suffer from poor pharmacokinetics and thiol reactivity. By targeting allosteric sites, specifically the G85 and alpha pockets, we have identified compounds that functionally disrupt the enzyme’s protective capacity without the need for reactive warheads. The high synergistic correlation (r = 0.81) between the G85 site and the catalytic centre suggests a potent regulatory mechanism where allosteric binding likely induces a conformational shift in the α3-helix, obstructing substrate access to the peroxidatic cysteine (C_p_). Furthermore, our spatial transcriptomics data demonstrates that PRDX5 expression is concentrated in the hypoxic tumour core rather than the invasive border, suggesting that malignant cells upregulate this pathway specifically to survive the metabolic stress of the core microenvironment.

Nevertheless, translating these biochemical hits into clinical efficacy requires addressing several key limitations. While we have shown that MPM cells exhibit PRDX5 dependent resistance to peroxynitrite and oxygen peroxide, the potential for redundant antioxidant systems, such as the glutathione-peroxidase axis or other PRDXs, to compensate for PRDX5 inhibition must be explored in more complex validation models^15,16,18,22^. Additionally, while our biochemical screen yielded an 8% hit rate, the *in vivo* stability and blood-brain barrier permeability (relevant for MPM metastasis) of these azetidine containing scaffolds remain unknown^31^. The immediate trajectory for this work involves the structural validation of our lead candidates co-crystallised with PRDX5 to provide definitive structural evidence of binding within the predicted allosteric pockets.

We speculate that PRDX5 inhibition could serve as a powerful sensitising strategy. By stripping epithelioid MPM cells of their specialised RNS and ROS detoxification machinery, we may significantly lower the threshold for cell death when combined with ROS generating standard of care chemotherapies or emerging therapies like cold physical plasma^32^. Ultimately, this study provides a template for precision oncology in mesothelioma, moving beyond “one-size-fits-all” treatments toward mechanistically driven interventions for the patients with the highest unmet need.

## Supporting information

Supplementary Information - PRDX5 biochemical screen results

## Acknowledgements

This study also makes use of data generated by the MOSAIC consortium (Owkin; Charité – Universitätsmedizin Berlin (DE); Lausanne University Hospital - CHUV (CH); Universitätsklinikum Erlangen (DE); Institut Gustave Roussy (FR); University of Pittsburgh (USA)), a non-interventional clinical trial registered under NCT06625203. We are grateful to patients, physicians, nurses and research assistants involved in the study.

We thank Sandrine Delbary Gossart, Andrew J. Pierce and Rita Santos for their scientific contributions and discussions on the study.

## Author contributions

JM, AMK, YB, LH, NJ, AJC, RC, AR, ER and EP contributed to the identification of PRDX5 as a target in MPM via the analysis of the MOSAIC consortium data. The co-authors from the MOSAIC centers contributed to the recruitment of patients, documentation of clinical data and generation of multi-omics data. JM, FM, AMK, EP, LP, VL, P-BA, CP and SG contributed to the target validation activities. GW, GY, QS, LS and JZ contributed to the identification of the allosteric pockets and conducted the virtual compound screen. JM, FM, NJ, CJ, AMK, LP, P-JZ, JS, SG and EP contributed to the development and performance of the biochemical screening assay and hit assessment.

## Materials and Methods

### Data Analysis

#### Data description

MOSAIC (Multi-Omics Spatial Atlas in Cancer), a multi-centre clinical Omics study is an ongoing initiative led by Owkin and 5 hospitals aiming to generate the world’s largest spatial atlas in cancer^23^. It collects and generates six data modalities (extensive clinical data, Hematoxylin and Eosin stained (H&E) slides, 10X Visium spatial transcriptomics, single-nuclei transcriptomics, bulk RNAseq, and whole-exome sequencing). In particular, the technologies used are Visium V2 Cytassist, applied to Formalin-fixed paraffin-embedded (FFPE) samples, and the captured area covers 6.5 mm x 6.5 mm; and Chromium Single Cell.

Here we analysed baseline mesothelioma samples from three centres UKER, CHUV and CR (Table 5).

**Table 5.** Summary of MOSAIC MPM samples available for the study.

| MOSAIC Cohort |  | UKER | CHUV | CR | Total |
| --- | --- | --- | --- | --- | --- |
| Total samples |  | 24 | 26 | 24 | 74 |
| Total samples per modality | Single-cell samples | 24 | 26 | 24 | 74 |
|  | Single-cell samples with histological annotation | 21 | 26 | 17 | 64 |
|  | Visium samples | 14 | 23 | 17 | 54 |

#### Availability of data and materials

The MOSAIC dataset supporting the conclusions of this article is not publicly available but access to a subset of 60 patients can be requested through the MOSAIC - Window (https://www.mosaic-research.com/mosaic-window). Specifically, 9 representative MPM patients used in this study are available upon request from EGA with accession ID EGAC50000000398.

#### Statistical methods and analysis

All analyses were performed in Python (v3.11). All tests were two-sided. Multiple-testing correction used Benjamini–Hochberg FDR within each analysis family.

#### Expression across single-cell data

Gene expression (*PRDX5, PRDX3*) was normalised using CPM and log-transform. For each sample and cell type, we reported (i) percentage of cells with expression > 0 and (ii) mean expression among expressing cells.

#### Association of gene signatures with *PRDX5/3* expression

Three signatures (hypoxia, ROS detoxification, RNS metabolism) were scored as the mean of member genes (excluding PRDX5 from gene sets when relevant). These signatures were picked either from the REACTOME or Gene Ontology datasets.

In malignant cells, cells were stratified into PRDX5-positive (logCPM > 0) versus PRDX5-negative (logCPM = 0). For each sample, we computed the median signature score in PRDX5-positive and PRDX5-negative cells and their paired difference (median in PRDX5-positive minus median in PRDX5-negative). Across samples, this paired difference was tested against zero using a Wilcoxon signed-rank test and FDR-adjusted across the three signatures. Specificity was assessed using a matched-bins gene-set null model: random gene sets of equal size were sampled to match the expression-bin composition of the observed signature, and an empirical two-sided p-value was computed from the null distribution of the median paired difference.

Association of gene signatures and PRDX5/3 expression with tumour regions Within Visium slides (Cancer Core), three spatial regions were defined: Tumour Islet (centre), TI Edge (margin band), and Stroma. Tumour spots were identified using a threshold on the malignant deconvolution fraction. Because slide signal-to-noise varies, the tumour-fraction threshold and morphological refinement parameters were selected per slide. Tumour islets were then refined by smoothing and connecting tumour spots, filling small holes, and removing small, isolated regions; the TI Edge was defined as a fixed-width band around the islet boundary.

Spot-level expression was computed as logCPM (target sum 1e6, log1p). PRDX5/PRDX3 and signature scores were summarised per slide by medians in Tumour Islet versus TI Edge, and the paired difference (median in TI Edge minus median in Tumour Islet) was tested across slides using a Wilcoxon signed-rank test with FDR adjustment across the three signatures. A matched-bins gene-set null model was computed analogously to the single-nuclei analysis.

### Detection of total PRDX5 protein in human tumours

#### Tissue Samples

Tissue samples of 4 µm sections from mesothelioma patients were acquired from the CRB (Centre de Ressources Biologiques) of Toulouse, France, in accordance with institutional ethical guidelines and approvals.

#### Tissue Preparation and Antigen Retrieval

After deparaffinisation; antigen retrieval was performed using CC1 buffer (Ventana Systems) for 40 minutes at 90°C in a Discovery ULTRA automated staining system.

#### Immunohistochemistry

Immunohistochemical staining for Peroxiredoxin 5 (PRDX5) was performed using a rabbit polyclonal antibody (Bio-Techne, catalogue number NBP3-32740, lot H680063042; stock concentration 1 mg/mL). The antibody was diluted 1:10,000 in antibody diluent and incubated for 60 minutes at 37°C. Detection was carried out using the OmniMap anti-Rb HRP Detection Kit (Ventana) following the manufacturer’s protocol. Slides were counter-stained with hematoxylin, dehydrated, and coverslipped.

#### Cell lines

The human malignant pleural mesothelioma (MPM) cell lines IST-MES-1, and IST-MES-2 were obtained from Creative Bioarray. MSTO-211H and the normal mesothelial control cell line MeT-5A were sourced from ATCC and Addexbio, respectively.

IST-MES-1 and IST-MES-2 cells were cultured in Dulbecco’s Modified Eagle Medium (DMEM) supplemented with 2 mM L-glutamine, 1% non-essential amino acids, and 10% fetal bovine serum (FBS). MSTO-211H cells were maintained in ATCC-formulated RPMI-1640 medium supplemented with 10% FBS. MeT-5A cells were cultured in Addexbio-formulated Medium 199/MeT-5A, enriched with 3.3 nM human recombinant epidermal growth factor (EGF), 870 nM zinc-free bovine insulin, and 10% FBS.

#### Detection of total PRDX5 protein in cancer cell lysates

Human MPM cell lines (IST-MES-1, IST-MES-2, MSTO-211H) and the normal mesothelial control cell line (MeT-5A) were detached, seeded, and cultured under standard conditions. After three days, when cells reached approximately 80% confluence, they were washed with phosphate-buffered saline (PBS) and lysed using RIPA buffer (Thermo Scientific, catalogue no. 89900) supplemented with protease and phosphatase inhibitors for protein extraction. Protein concentrations were determined using the bicinchoninic acid (BCA) assay. Equal amounts of protein were then analysed using the JESS capillary-based protein detection system (ProteinSimple) according to the manufacturer’s instructions. Antibody dilutions were optimised for detection as follows: PRDX5 (Proteintech, #67599-1-Ig) at 1:50 and PRDX3 (Proteintech, #10664-1-AP) at 1:25. Data were normalised using the Simple Western Total Protein Assay (performed on the same run).

#### Cytotoxicity measurement

Cells were seeded at the following densities in 96 well plates: IST-MES-1 (5,000 cell per well), IST-MES-2 (3,000 cell per well), MSTO-211H (6,000 cell per well)) and MeT-5A (1,500 cell per well). After 24h (to allow cell attachment), cells were treated with an exogeneous concentration range of Peroxynitrite (250-4000µM) or H_2_O_2_ (5-640 µM). After 5 days of treatment, cell viability was assessed using CellTiter-Glo measurement (luminescence value).

### Identification of putative allosteric pockets

#### Structure Preparation

The atomic coordinates of the seven PRDX5 crystal structures were retrieved from the RCSB PDB^33^. Co-crystallised water molecules, glycerol, dimethyl sulfoxide, ions, and substrate ligands were removed, retaining only the protein structures. Preparation was performed using MOE (version 2024.06) with the integrated Structure Preparation module. The structures were optimised and protonated using Protonate3D at a physiological pH of 7.4 under the Amber14:EHT force field^34^. All structures were aligned to the reference structure 3MNG within the MOE environment. The pairwise RMSD values between the structures were calculated using PyMOL (version 2.5.7)^35^, and the resulting matrix is presented as a heatmap in Supplementary Figure S2.

#### Site Prediction

The seven prepared structures were submitted for AlloDeep calculations with the designated substrate site. Predicted allosteric sites were subsequently clustered based on their spatial overlap. The apo structure 2VL3 served as the starting configuration for MD simulation. The MD simulations were conducted for a total of 600 ns using the APSF force field with Amber (version 22)^36^, followed by AlloTango analysis to evaluate site synergy. All predicted sites were inspected in PyMOL to confirm their geometry and precise localisation, ensuring logical clustering. Site-specific properties were integrated based on cluster assignments and subjected to statistical characterisation.

### Virtual Screening

#### Compound Library Filtering and Preparation

To ensure library quality and minimise false positives, a multi-step filtering protocol was implemented. The initial Enamine Screening Collection was subjected to structural filters using RDKit (version 2019.09)^37^. Molecules were filtered via PAINS and REOS rules to exclude compounds with reactive, toxic, or undesirable functional groups. Additionally, compounds were filtered by molecular weight (150 < MW < 500 Da). The remaining high-quality molecules were prepared in MOE using the Wash module, which included adding explicit hydrogens and assigning ionisation states at pH 7.4. Three-dimensional (3D) conformations were generated and energy-minimised using the Amber14:EHT force field. Finally, the molecules were converted into PDBQT format using the AutoDockTools (ADT) suite (version 1.5.7)^38^, with Gasteiger partial charges assigned and non-polar hydrogens merged.

#### Molecular Docking

High-throughput molecular docking was performed using smina (version 2019.10.15), a fork of AutoDock Vina^39^ that focuses on improving scoring and minimisation. Receptor structures were prepared in ADT by assigning Gasteiger charges and converting the pre-protonated proteins into PDBQT format. All protein residues were kept rigid during the simulations. The docking search space was defined by a cubic grid box (20 × 20 × 20 Å3) centred on the Cartesian coordinates identified by AlloDeep. The default smina scoring function was employed, with the exhaustiveness set to 8 and num_modes limited to 1 to balance conformational sampling with computational throughput. Docking was executed on a high-performance computing (HPC) cluster using multi-threaded parallelisation.

#### Chemical Space Clustering

To analyse structural diversity, molecules were represented as Morgan fingerprints^40^ (ECFP4 equivalent) using RDKit, with a radius of 2 and a bit-length of 2048. The resulting vectors were converted into a float32 matrix for efficiency. Clustering was performed using Faiss (Facebook AI Similarity Search)^41^, employing a K-means algorithm to partition the library into 100,000 clusters based on Euclidean distance. Centroids were calculated using the IndexFlatL2 index with a maximum of 50 iterations. Medoids (molecules closest to each centroid) were selected as representative scaffolds to reduce redundancy while preserving chemical diversity.

#### Post-Docking Analysis

Compounds were ranked by their predicted binding affinity (smina score, kcal/mol). The top-scoring candidates were further filtered based on their internal strain energy and PoseFit score. The internal strain energy was assessed in RDKit using the MMFF94s force field^42^. Pose validity was evaluated using PoseFit, which measures the similarity of interaction fingerprints with experimental data in the PDB. The remaining high-priority compounds were clustered into 300 groups for each site. Finally, the binding modes were visually inspected in MOE to evaluate critical interactions, including hydrogen bonds, salt bridges, and hydrophobic contacts with key residues. During the manual curation process, compounds possessing structural alerts or substructures generally deemed unfavourable in medicinal chemistry were excluded.

### Protein expression and purification

Human PRDX5 (54-214) wild type was cloned with an N-terminal 8xHIS tag followed by a TEV protease site and an Avi tag for biotinylation of protein in pET29a for expression E. coli. Similarly, human TrxR and Trx genes were cloned in pET29a with an N-term 8xHIS tag for the biochemical assays. The genes were expressed in the Rosetta (DE3) pLysS strain of E. coli by induction with 0.5 mM IPTG at a cell density of 0.8 and cell were grown at 18°C overnight. The cells were harvested and lysed by sonication in lysis buffer containing 20 mM sodium phosphate (pH 7.4), 500 mM NaCl, 20 mM Imidazole, 2 mM DTT, followed by elution of protein in the lysis buffer containing 500 mM Imidazole. The protein was further purified by size-exclusion chromatography on a HiLoad Superdex 75 26/600 GL in a buffer containing PBS pH7.4 2mM DTT. The dimeric peak was collected, concentrated and aliquoted for assays. The proteins were observed to be of >95% purity and were confirmed by intact MS.

### PRDX5 activity assay (Thioredoxin assay)

Enzymatic activity of PRDX5 was measured using the thioredoxin system. The thioredoxin assay was adapted from previously described protocol^30^ become high throughput screening-compatible assay to identify and characterise/profiling of small molecule inhibitors. The assay was performed in a 8 µL reaction mixture, containing Phosphate buffer saline (pH = 7.1 Gibco), 250 µM NADPH, 760 nM human TrxR, 11 µM human Trx, 1µM WT Prdx5 and 500µM H_2_O_2_. First, PRDX5 protein was dispensed on compounds already in a well of 1,536 well plate (Corning 3724). The mix was incubated at 37 °C for 30 min. Then Trx, TrxR and NAPDH were dispensed on WT PRDX5 protein, and the reaction was initiated by the addition of 500 µM H_2_O_2_. The mix was incubated at 37 °C for 1 h 15 min before reading fluorescence at 460 nm using Pherastar device (BMG Labtech). Results were analysed in GraphPad prism software using median and standard deviation.

### Screening data analysis

Thioredoxin assay was performed in triplicate. Results were analysed on Genedata Screener software. Quality controls were validated for each plate using RZ’, signal to background and auranofin positive control IC_50_. Raw fluorescence intensity signals are normalised by 0-100% activity using DMSO and auranofin at 30 µM controls, respectively. The fit model used was Smart Fit condensing. It identifies and excludes outliers and selects the best fit model, Hill fit or constant fit. The data mode (decreasing, increasing or inactive) is set to follow scale reference (based on controls). To rank compounds, we considered the data mode of the curve, relative potency qIC_50_ and maximal activity (median of replicates at maximal tested concentration).

## Supplementary Materials

**Supplementary Figure S1:**
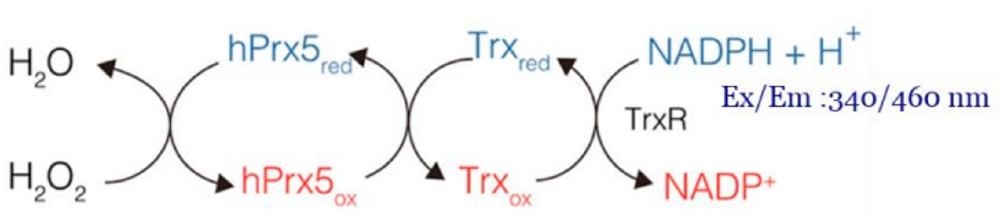
The biochemical reaction used to detect inhibition of PRDX5 in the screening assay.

**Supplementary Figure S2:**
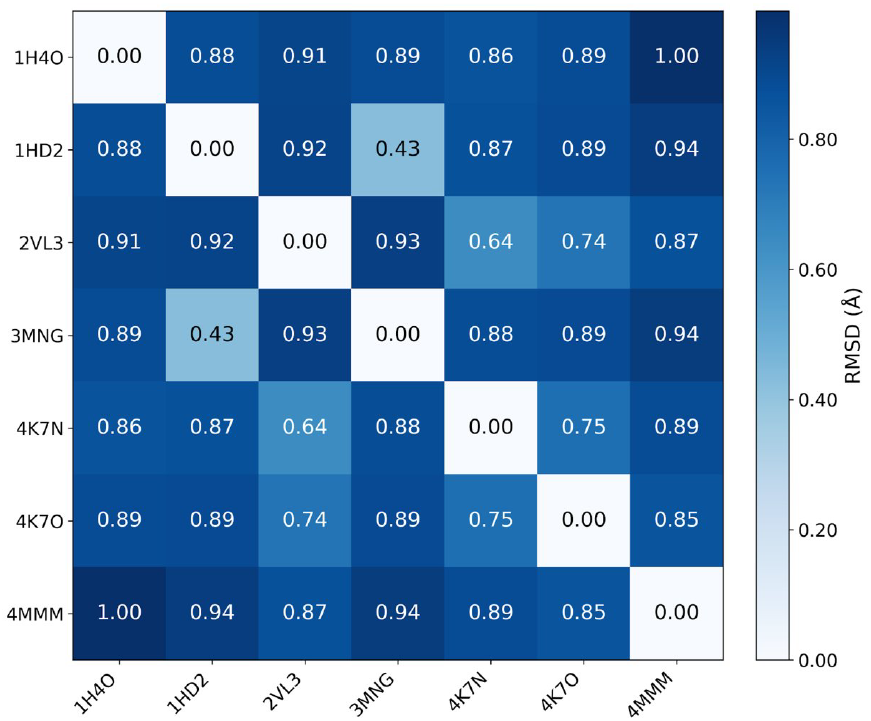
Matrix of pairwise distances (RMSD) between all PRDX5 crystal structures.

**Supplementary Figure S3:**
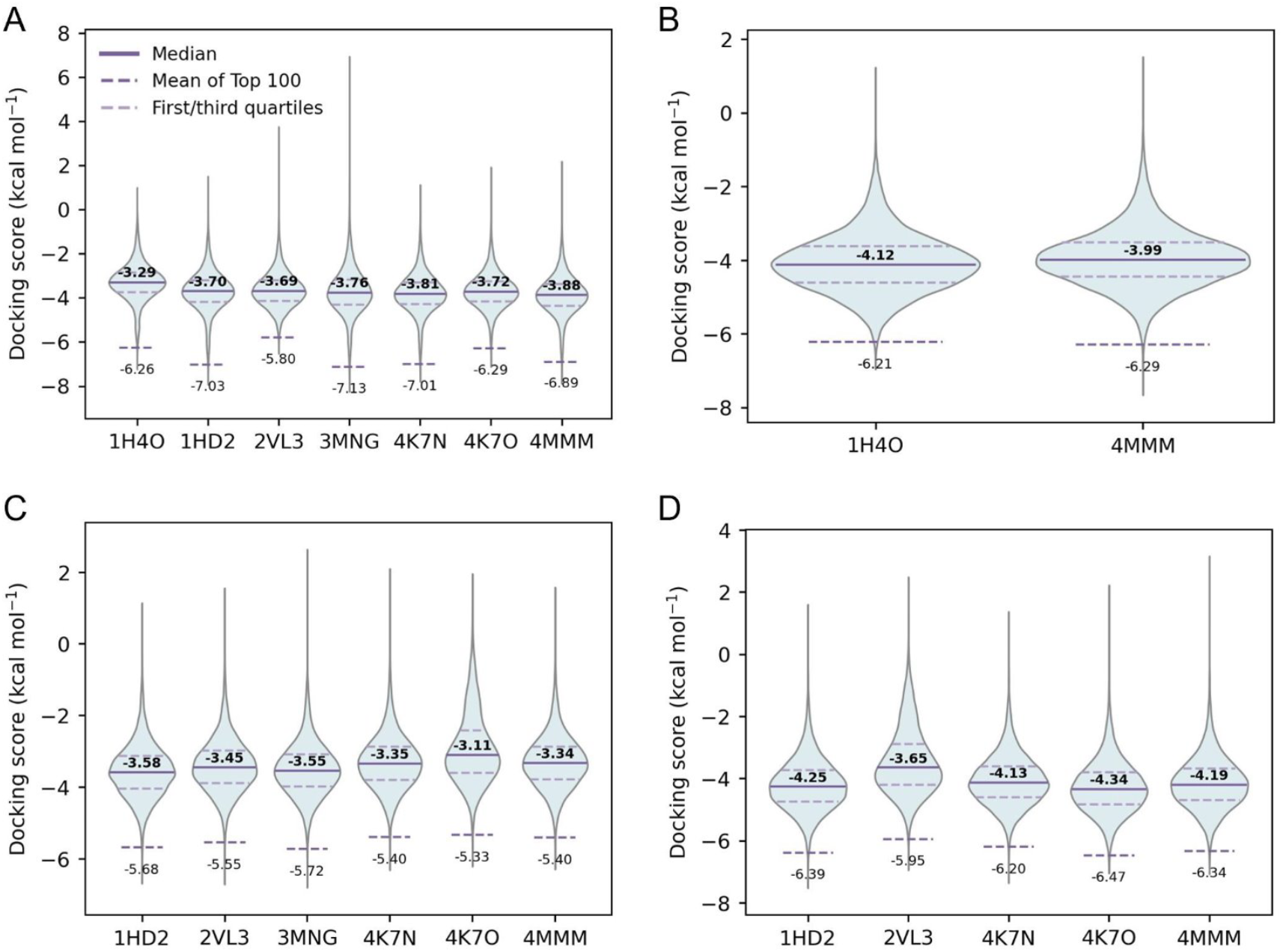
Docking score distributions for various pocket conformations at the (A) substrate site, (B) G85 site, (C) Alpha3 site and (D) Alpha5 site.

### Supplementary Note

We additionally share a .csv file with the chemical structures of the 452 tested compounds. We describe the columns here:

- SMILES: the SMILES representation of the molecule
- Name_Enamine: the molecule ID
- Site: the binding site corresponding to the virtual hit
- Occurrence of Compound within the Top300 across different conformations: number of starting protein structures for which the compound was ranked in the top 300 molecules
- SiteConformation: PDB id for the corresponding protein structure
- Smina scores: docking score from Smina
- LigandEfficiency: A normalised metric used to compare the binding affinities of compounds across different molecular sizes. It is calculated as the absolute docking score divided by the number of non-hydrogen (heavy) atoms. By penalising excessive molecular weight, LE helps prioritise compounds with optimal drug-like properties. Units: kcal/mol per heavy atom.
- PoseFit score: This score evaluates pose validity by comparing the interaction fingerprints of docked poses against conserved fragment-binding patterns derived from the PDB database. Ranging from 0 to 1, higher scores indicate superior alignment with experimentally validated interaction motifs, suggesting a more biologically plausible binding mode.
- StrainEnergy_MMFF94s: Defined as the difference in total potential energy between the docked conformation and its energy-minimised state, calculated using the MMFF94s force field via RDKit. High values represent a significant energetic penalty for the conformational adaptation required for binding, which may decrease overall binding probability. Units: kcal/mol.
- HBD: H-bond donors
- HBA: H-bond acceptors
- Lead-likeness: Binary predicted Lead-likeness from MW, logD, ring count, rotatable bond count, HD count and HA count
- Bioavailability: Binary predicted bioavailability from MW, logP, PSA, fused aromatic ring count, rotatable bond count, HD count and HA count
- Potential_Covalent_Motif: number of chemical moieties that could be involved with covalent bonding
- Severity_Score: Severity score for Evotec structural alert flags
- EvoAlert: Evotec structural alert flags
- PRIM_DataMode: Activity mode of the compound: either increasing, increasing (weakly active) or inactive
- PRIM_operator: operator for PRIM_Potency.uM
- PRIM_Potency.uM: Calculated potency in µM for increasing compounds
- PRIM_Sinf: Top plateau set for curve fitting
- PRIM_nHill: Hill slope
- PRIM_Activity (cmax): Activity % at the max concentration
- PRIM_pIC50: pIC50: –log10(IC50 in molar)
- PRIM_LE: Ligand Efficiency: (1.37*pIC50)/N heavy atoms
- PRIM_Lipe: Lipophilic Efficiency: LipE=pIC50−logD(pH7.4)
- PRIM_Lipe_LogP: Lipophilic Efficiency: LipE=pIC50−logP
- HitComp_Purity: Purity of compound by LC/MS QC

